# Females are protected from semaglutide-induced muscle loss in *ob/ob* mice

**DOI:** 10.64898/2026.03.03.709376

**Authors:** Subhasmita Rout, Takuya Karasawa, Shinya Watanabe, Amandine Chaix, Micah J Drummond, Katsuhiko Funai, Ran Hee Choi

## Abstract

Obesity is a major contributor to cardiometabolic disease, and pharmacological therapies such as semaglutide are increasingly used to induce weight loss. However, the commonly used diet-induced obesity model in C57BL/6J mice is limited by relative resistance to weight gain in females, complicating the study of sex-specific effects. Here, we used leptin-deficient *ob/ob* mice, which develop severe early-onset obesity in both sexes, to investigate sex-specific responses to semaglutide on skeletal muscle mass, function, and mitochondrial metabolism. The *ob/ob* mice were treated daily with semaglutide or vehicle for three weeks, followed by assessments of body composition, muscle and organ mass, muscle contractile function, and mitochondrial efficiency. Semaglutide induced comparable reductions in body weight and food intake in both sexes but elicited distinct sex-specific changes in body composition. Male mice exhibited losses in both skeletal muscle and organ mass, whereas female mice preferentially lost fat and organ mass while preserving skeletal muscle. Despite these divergent structural adaptations, muscle force generation remained intact in both sexes. Collectively, these findings reveal pronounced sexual dimorphism in skeletal muscle and metabolic remodeling during pharmacologically induced weight loss, highlighting the importance of considering biological sex when evaluating the metabolic and therapeutic effects of anti-obesity interventions.

**Article Highlight:**

- C57BL/6J mice are limited by relative resistance to weight gain in females, complicating the study of sex-specific effects. So, we wanted to determine the sex-specific effect of semaglutide on skeletal muscle function, and mitochondrial metabolism in *ob/ob* mice.
- We assessed body composition and ex-vivo muscle force following the treatment and found that the female *ob/ob* mice are protected from semaglutide-induced skeletal muscle mass loss.
- These findings demonstrate sex-specific effects of semaglutide, highlighting the need to consider biological sex in GLP-1RA–based therapies.

## Introduction

Obesity has reached alarming prevalence worldwide and is a principal contributor to type 2 diabetes, cardiometabolic diseases, and obesity-associated cancers (1, 2). Its accelerating trajectory represents a major public health challenge and a leading cause of premature mortality. Weight loss remains the most effective strategy to improve metabolic health, yet the physiological adaptations that accompany weight reduction, such as reduced energy expenditure and loss of lean mass, often challenge long-term weight maintenance (3–5). Pharmacological therapies for obesity management, including glucagon-like peptide-1 receptor agonists (GLP-1RAs) such as semaglutide, have emerged as powerful means to achieve and sustain clinically meaningful weight loss (10–15% over a year) compared with lifestyle intervention (5-10% over a year) (6–8). In addition to appetite suppression, the effects of GLP-1RAs on systemic metabolic and peripheral tissues, including skeletal muscle, adipose tissue, and liver, though the tissue-specific adaptations remain incompletely understood (9–11).

Skeletal muscle is a principal regulator of energy homeostasis, accounting for a majority of insulin-stimulated glucose disposal and a critical determinant of resting and activity-induced energy expenditure (12, 13). Across dietary, surgical, and pharmacologic weight loss interventions, reductions in fat mass are accompanied by loss of fat-free mass (FFM), of which skeletal muscle constitutes a substantial portion. In our previous work, semaglutide-induced weight loss significantly decreases FFM, yet the decline was disproportionate to losses in individual hindlimb muscle masses (14). Additionally, this FFM loss lowers resting energy expenditure and predicts weight regain (4, 5). Part of this adaptive response reflects a lower mitochondrial fuel cost for ATP synthesis, such as increased coupling efficiency, which reduces whole-body energy expenditure during the weight reduced state (15, 16). We also observed that weight loss induced by semaglutide, or dietary intervention improves skeletal muscle mitochondrial oxidative phosphorylation (OXPHOS) efficiency in diet-induced obese (DIO) mice, consistent with tighter ATP to O_2_ (P/O) coupling (15, 17). These observations underscore that skeletal muscle plays a central role in the energetic consequences of weight loss and motivate testing how semaglutide modulates muscle mass, function, and mitochondrial efficiency under different contexts than leptin-intact DIO mice.

DIO is the most prevalent form of obesity in humans and the most widely used experimental model for studying weight loss. However, female C57BL/6J mice are relatively resistant to diet-induced weight gain, limiting their utility for weight loss studies in females (18, 19). In contrast, the leptin-deficient *ob/ob* mouse represents a distinct genetic model of obesity characterized by hyperphagia, early onset, and severe adiposity resulting from a mutation in the Lep gene. This mutation leads to the absence of functional leptin, causing profound obesity, hyperphagia, and metabolic dysregulation beginning early in development (21). Consequently, the *ob/ob* model provides a robust experimental framework to investigate semaglutide-induced weight loss in both male and female mice.

In the present study, we aimed to investigate the effect semaglutide-induced weight loss on muscle force producing capacity and mitochondrial efficiency in *ob/ob* mice. We hypothesized that semaglutide-induced weight loss in *ob/ob* mice produces sex-specific alterations in lean mass and contractile force during weight reduction.

## Materials and methods

### Animals

Eight-week-old male and female *ob/ob* mice (IMSR_JAX:000632) were purchased from Jackson Laboratory. Upon arrival, mice were housed in a temperature-controlled room (22°C) with a 12-hour light/dark cycle and provided with a standard chow diet and deionized water *ad libitum*. Mice were allowed to acclimate for a period of 4 weeks. Body mass and food consumption were measured weekly during this acclimation period until mice reached approximately 50g. Following the 4-week acclimation period, mice were randomly divided into two experimental groups where one group received daily subcutaneous injections of 3 nmol/kg body mass semaglutide (Novo Nordisk), and another group received daily subcutaneous injections of an equivalent volume of vehicle (phosphate-buffered saline, PBS). Injections were administered daily for a period of 3 weeks and during the treatment period, body mass and food consumption were measured daily. At 15 weeks of age, before the termination the mice were fasted for 2 hours followed by intraperitoneal injection of 80 mg/kg ketamine and 10 mg/kg xylazine to proceed for the dissection. The muscle tissues, including tibialis anterior (TA), extensor digitorum longus (EDL), plantaris (PLA), soleus, gastrocnemius, and quadricep were harvested, snap-frozen in liquid nitrogen, and stored at -80°C for subsequent biochemical and molecular analyses. All animal experiments were performed with the approval of the Institutional Animal Care and Use Committees at the University of Utah.

### Body composition

Whole-body fat and lean mass were determined using quantitative nuclear magnetic resonance (NMR) spectroscopy (Minispec, Bruker) on 8 weeks of age and again on 12 weeks of just before the treatment was started. The body composition was again measured after 1 week and 3 weeks of post treatment which was 13 weeks and 15 weeks of age.

### Ex vivo muscle force production

Muscle force capacity was assessed using an *ex vivo* system (Model 801C, Aurora Scientific, Aurora, Canada). The EDL and soleus muscles were freshly harvested and then securely fastened at their tendons. Each muscle was individually mounted within a tissue bath, attached to an anchor at one end and a force transducer at the other. The bath contained continuously oxygenated Krebs-Henseleit buffer (KHB) maintained at 37°C. Once the optimal length was established, muscles were incubated in fresh KHB for a 5-minute equilibration period, following this, force-frequency was initiated. Muscles were stimulated at progressively increasing frequencies (10, 20, 30, 40, 60, 80, 100, 125, 150, and 200 Hz), with a 2-minute rest interval between each stimulation frequency. After the force measurements, muscle length and mass were precisely measured to calculate the cross-sectional area (CSA). Force production data were analyzed using Aurora Scientific’s DMAv5.321 software. Results are reported as specific force, which is calculated by normalizing the absolute force to the muscle’s CSA.

### Tissue sectioning and histology

Freshly harvested TA muscles were embedded in optimal cutting temperature media and were snap freeze in liquid nitrogen immediately and stored in -80°C freezer. Muscle tissues were sectioned at 8 μm thickness with the Leica CM1860 cryostat (Leica Biosystems) where the Cryostat chamber was maintained with a temperature of -20°C during the period. The tissue sections were collected on a positively charged coated slide and stored in -20°C for histology.

To determine the fiber type in the TA muscle, the muscle sections were retrieved from the -20°C and allowed to incubate at room temperature for 15 min. Muscle sections were prefixed with 10% paraformaldehyde for 10 minutes followed by 1X PBS washing. Then sections were incubated with Mouse-on-Mouse blocking reagent for 1 hour at room temperature (RT) to block the endogenous immunoglobulins following overnight at 4°C incubation with specific primary antibodies targeting different myosin heavy chain (MHC) isoforms: MHC IIa (DHSB, SC-71) and MHC IIb (DHSB, BF-F3) with 2.5% goat serum in 1X PBS. On the next day, muscle tissue sections were incubated with anti-mouse secondary antibody IgG2b (A 21242), IgG1 (A 21120) and IgM (A 21426) and wheat germ agglutinin (WGA) Alexa Fluor 488 conjugate (Thermo Fisher Scientific, 0.004 mg/mL) with 2.5% goat serum in 1X PBS for 1 hour at RT followed by PBS wash. Muscle sections were mounted with mounting medium (Vector Laboratories, H-1000) and imaged using Zeiss Axioscan 7 Slide Scanner (Carl Zeiss). The muscle fiber typing was analyzed using cellpose-SAM model as previously described (20).

### Permeabilized muscle fiber bundles (PmFB)

Permeabilization of red gastrocnemius muscle fibers was performed as previously described in (17). Briefly, a small portion of red gastrocnemius muscle was separated and incubated in buffer X [7.23mM K_2_EGTA, 2.77 mM CaK_2_EGTA, 20 mM imidazole, 20 mM taurine, 5.7 mM ATP, 14.3 mM phosphocreatine, 6.56 mM MgCl_2_·6H_2_O, and 50 mM 2-(N-Morpholino) ethane sulfonic acid potassium salt (K-MES) (pH7.4)]. Muscle tissue was teased in buffer X to separate individual muscle bundles and permeabilized by incubation in saponin (30 μg/ml) for 30 minutes at 4°C. Following permeabilization, the muscle fiber bundles were washed with buffer Z [105 mM K-MES, 30 mM KCl, 10 mM K_2_HPO_4_, 5 mM MgCl_2_·6H_2_O, 0.5 mg/mL BSA, and 1 mM EGTA, pH 7.4] with 0.5 mM pyruvate and 0.2 mM malate for 15 min at 4°C. Lastly, the muscle fiber bundles were placed in buffer Z until the experiment.

### High-resolution respirometry and fluorometry

Oxygen consumption in PmFB was measured using the Oroboros Oxygraph-2K (Oroboros Instruments) as previously described (17). ATP production was determined by enzymatically coupling ATP production to NADPH synthesis, with the reaction monitored using a Horiba Fluorolog-QM (Horiba Scientific) as previously described (21). Both experiments were conducted in buffer Z, supplemented with 20 mM creatine monohydrate. To inhibit myosin adenosine triphosphatase, 10 μM blebbistatin was added to the buffer. The experiments were initiated with the sequential addition of substrates: 0.5 mM malate, 5 mM pyruvate, 5 mM glutamate, and 10 mM succinate. Respiration and ATP production were recorded with subsequent stepwise titrations of ADP at concentrations of 20, 200, and 2000 μM.

### Western blot

The frozen whole gastrocnemius muscle (∼50mg) homogenized in ice-cold RIPA buffer (ThermoFisher Scientific, 89901) supplemented with a protease inhibitor cocktail (ThermoFisher Scientific, 78446). Protein concentration was determined using a BCA protein assay kit (ThermoFisher Scientific, 23225). For protein analysis, equal amounts of protein were separated by electrophoresis on a 4-20% gradient SDS-polyacrylamide gel (Bio-Rad). The proteins were then transferred to polyvinylidene fluoride membranes (ThermoFisher Scientific, 88518). The membranes were blocked for 1 hour at room temperature in 5% skim milk in Tris-buffered saline with 0.1% Tween-20 (TBST). Following the blocking step, the membranes were incubated with primary antibodies against oxidative phosphorylation (OXPHOS) complexes (Abcam, MS604-300) and citrate synthase (Abcam, ab96600) at 4°C overnight. The next day, the membranes were washed with TBST 3 times for 5 minutes each and incubated with an anti-mouse secondary antibody (1:5000 dilution) in 5% skim milk for 1 hour at room temperature. After another 3 times of washes with TBST, the membranes were incubated with SuperSignal™ West Pico PLUS Chemiluminescent Substrate (Thermo Scientific, 34580). The resulting chemiluminescence was imaged using a ChemiDoc Imaging System (Bio-Rad) and quantified with Image Lab Software (Bio-Rad).

### Statistical analysis

GraphPad Prism 10.4.1 software was used to do statistical analysis such as unpaired Student’s t-test to compare the lean tissue difference and two ways of variance (ANOVA) to analyze the body weight and food consumption in semaglutide treated and vehicle groups in both male and female mice. Additionally, to analyze the muscle force-frequency change between both the groups in both the sexes, again two ways ANOVA was performed. All data are represented as mean ± SEM, and statistical significance was set at p ≤ 0.05.

## Results

### Semaglutide-induced weight loss results from reduced food consumption in *ob/ob* mice

Eight-week-old male and female *ob/ob* mice were maintained on a standard chow diet for four weeks where their average body weight reached ∼50g and ∼52g for male and female respectively, after which they were treated with either 3 nmol/kg semaglutide or vehicle for three weeks (Figure 1A) and the food and body weight recorded daily. Semaglutide-treated mice exhibited a ∼15% reduction in food intake in males and ∼18% in females. Prior to treatment, weekly food intake averaged ∼24 g/week in males and ∼28 g/week in females. By the end of the treatment period, food intake decreased to ∼20 g/week in males and ∼23 g/week in females, remaining below baseline levels (Figure 1B). In contrast, vehicle-treated mice maintained consistent food consumption throughout the experimental period.

**Figure 1.**
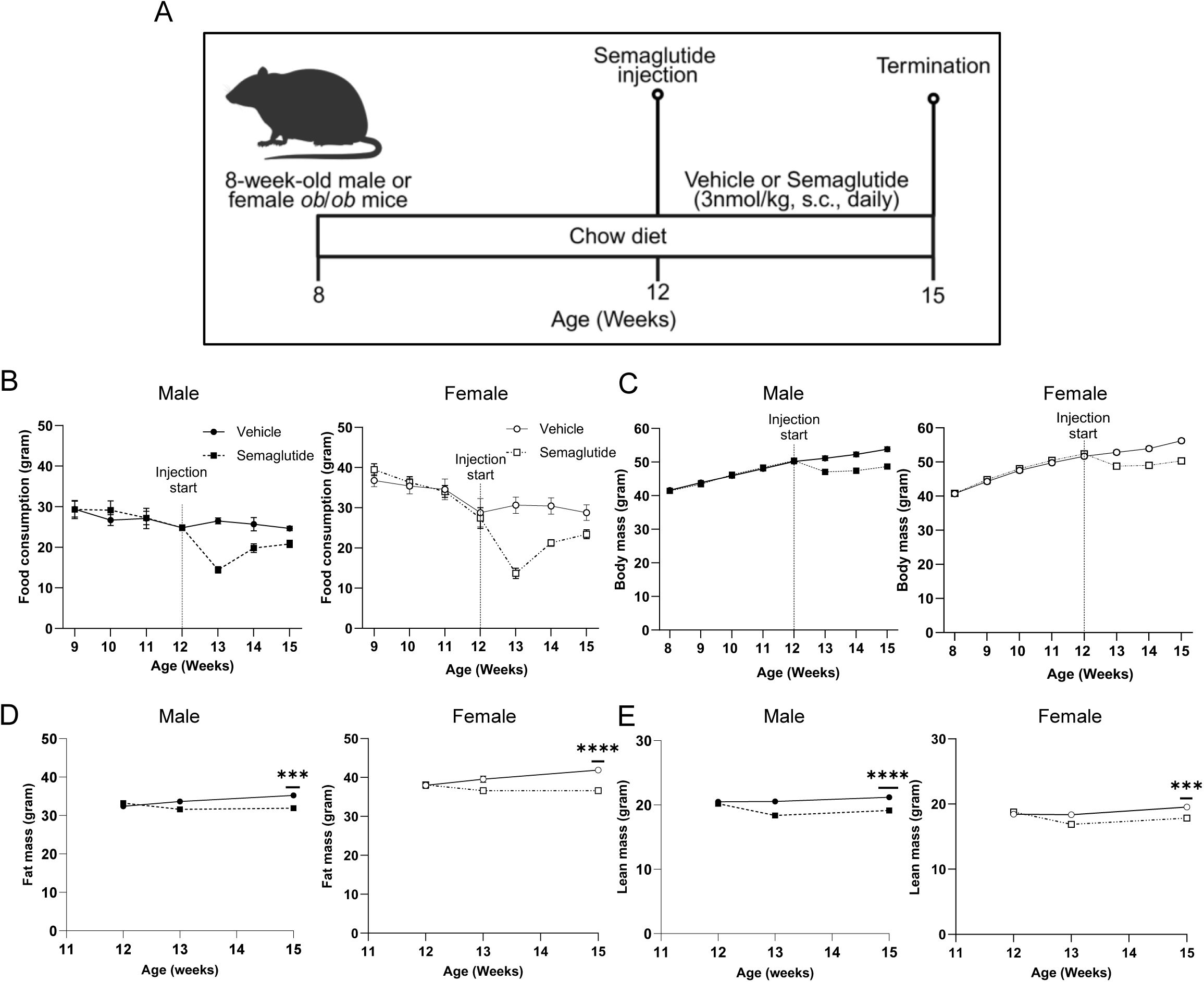
Semaglutide induced reduction in body mass, food intake and lean mass. A: Experimental setup showing 8-week-old *ob/ob* mice were maintained on chow diet throughout the experiment. On the 12th week, semaglutide treatment was performed daily for 3 weeks till 15th week old. B: Food consumption in both male and female after semaglutide treatment compared to vehicle group. C: Body mass of both male and female after semaglutide treatment. D & E: Fat mass and lean mass was measured by NMR across the semaglutide injection period. Data is represented as mean ± SEM *** p<0.001 and **** p<0.0001 significant difference vs. corresponding controls. n=12/group for both male and female.

The reduction in food intake in semaglutide-treated mice was accompanied by a decline in body weight. After three weeks of treatment, body weights were ∼48 g in males and ∼50 g in females, whereas vehicle-treated mice showed a progressive increase in body weight, reaching ∼53 g (males) and ∼56 g (females) by the end of the study (Figure 1C). These differences corresponded to sustained body weight reductions of ∼9–10% relative to vehicle-treated controls (Table 1). Collectively, these findings suggest that the observed weight loss following semaglutide treatment was largely attributable to decreased food consumption.

**Table 1:** Individual muscle and organ tissue weight of both male and female semaglutide and vehicle treated ob/ob mice at 3 weeks.

|  | Male |  |  |  | Female |  |  |  |
| --- | --- | --- | --- | --- | --- | --- | --- | --- |
|  | Vehicle | Semaglutide | % difference | T-test (p-value) | Vehicle | Semaglutide | % difference | T-test (p-value) |
| Body mass | 53.8 ± 0.7 | 48.8 ± 0.5 | -9.2* | < 0.0001* | 56.2 ± 0.6 | 50.3 ± 0.7 | -10.5* | < 0.0001* |
| Fat mass | 35.2 ± 0.6 | 31.9 ± 0.4 | -9.5* | 0.0005* | 41.9 ± 0.5 | 36.6 ± 0.6 | -12.7* | < 0.0001* |
| Lean mass | 21.9 ± 0.3 | 19.1 ± 0.3 | -9.8* | 0.0001* | 19.5 ± 0.2 | 17.8 ± 0.3 | -8.6* | 0.0003* |
| i WAT | 4090.5 ± 176 | 3751.9 ± 100 | -8.3 | 0.1026 | 4391.1 ± 74 | 4090.7 ± 79 | -6.8* | 0.0123* |
| g WAT | 2860.2 ± 98 | 2880.5 ± 95 | -0.7 | 0.8843 | 4867.7 ± 196 | 3949 ± 177 | -18.9* | 0.0024* |
| EDL | 7.8 ± 0.2 | 7.3 ± 0.1 | -6.3* | 0.0503* | 7.1 ± 0.2 | 7 ± 0.1 | -1.4 | 0.6501 |
| Soleus | 7.8 ± 0.3 | 7.2 ± 0.2 | -7.8 | 0.0945 | 6.4 ± 0.1 | 6.5 ± 0.2 | 1.8 | 0.6048 |
| TA | 32 ± 0.6 | 29.5 ± 0.4 | -7.6* | 0.0048* | 27.4 ± 0.5 | 26.8 ± 0.5 | -1.9 | 0.4554 |
| Plantaris | 12 ± 0.3 | 11.6 ± 0.2 | -2.9 | 0.2993 | 9.4 ± 0.2 | 9.5 ± 0.2 | 0.7 | 0.8187 |
| Gastroc | 86.8 ± 1.9 | 80.7 ± 1.5 | -6.8* | 0.0259* | 68.8 ± 1.9 | 68.1 ± 1.5 | -1.0 | 0.7716 |
| Quadricep | 119.9 ± 4.5 | 108.1 ± 2.2 | -9.8* | 0.0278* | 98.1 ± 2.1 | 96.5 ± 2.9 | -1.6 | 0.6735 |
| Liver | 3538.1 ± 91 | 2250.2 ± 75 | -36.4* | < 0.0001* | 3827.2 ± 144 | 2149 ± 85 | -43.8* | < 0.0001* |
| Heart | 105.8 ± 2.8 | 93.8 ± 1.3 | -11.4* | 0.0007* | 103 ± 2.4 | 94.1 ± 0.7 | -8.7* | 0.0010* |
| Spleen | 63.9 ± 2 | 46.3 ± 2.2 | -27.6* | < 0.0001* | 55.8 ± 1.7 | 41.8 ± 1.2 | -25.2* | < 0.0001* |
| Lungs | 115.2 ± 4.4 | 106.6 ± 3.3 | -7.5 | 0.1283 | 112.8 ± 2.0 | 107.9 ± 3.2 | -4.4 | 0.2340 |
| Diaphragm | 65.5 ± 2.4 | 59.6 ± 2.3 | -9.1 | 0.0910 | 64.4 ± 1.6 | 60.5 ± 2.4 | -5.0 | 0.2104 |
| Kidney | 327.3 ± 8.2 | 296.6 ± 5.7 | -9.3* | 0.0052* | 327.3 ± 6.9 | 284.1 ± 7.6 | -13.2* | 0.0005* |

### Semaglutide-induced weight loss accompanied by the reduction in fat and lean mass in *ob/ob* mice

Semaglutide intervention has previously been associated with concurrent reductions in both fat and lean mass in humans and rodent models of obesity (22–24). To determine whether similar compositional changes occur in leptin-deficient *ob/ob* mice, characterized by adipocyte hyperplasia and abnormal lipid storage, we monitored body composition over the three-week treatment period. Three weeks of semaglutide treatment showed a significant reduction in fat mass and lean mass in both the sexes compared to vehicle treated mice (Figure 1D&E, Table 1). The reduction in fat mass was approximately 9.5% and 12.7% in male and female mice respectively (Table 1). Additionally, semaglutide treatment showed a dramatic reduction of lean mass ∼9.8% in male and ∼8.6% in female compared to vehicle treated mice (Table 1).

### Semaglutide treatment produces sex-dependent remodeling of adipose and muscle masses in *ob/ob* mice

Our previous study showed that semaglutide-induced weight loss was accompanied by reductions in both fat and lean mass; however, the decrease in total lean mass was not fully accounted for by proportional losses in individual skeletal muscles (14). In the present study, we tested whether semaglutide similarly alters tissue-specific mass distribution. Following three weeks of semaglutide treatment, individual skeletal muscles, inguinal white adipose tissue (iWAT), and gonadal white adipose tissue (gWAT) were harvested and weighed immediately. A clear sex-specific difference was observed in adipose tissue responses. Semaglutide-treated male *ob/ob* mice did not exhibit significant changes in either iWAT or gWAT mass compared with vehicle-treated controls (Figure 2A). In contrast, semaglutide-treated female *ob/ob* mice displayed significant reductions in both iWAT (∼6.8%) and gWAT (∼18.9%) (Figure 2B, Table 1).

**Figure 2.**
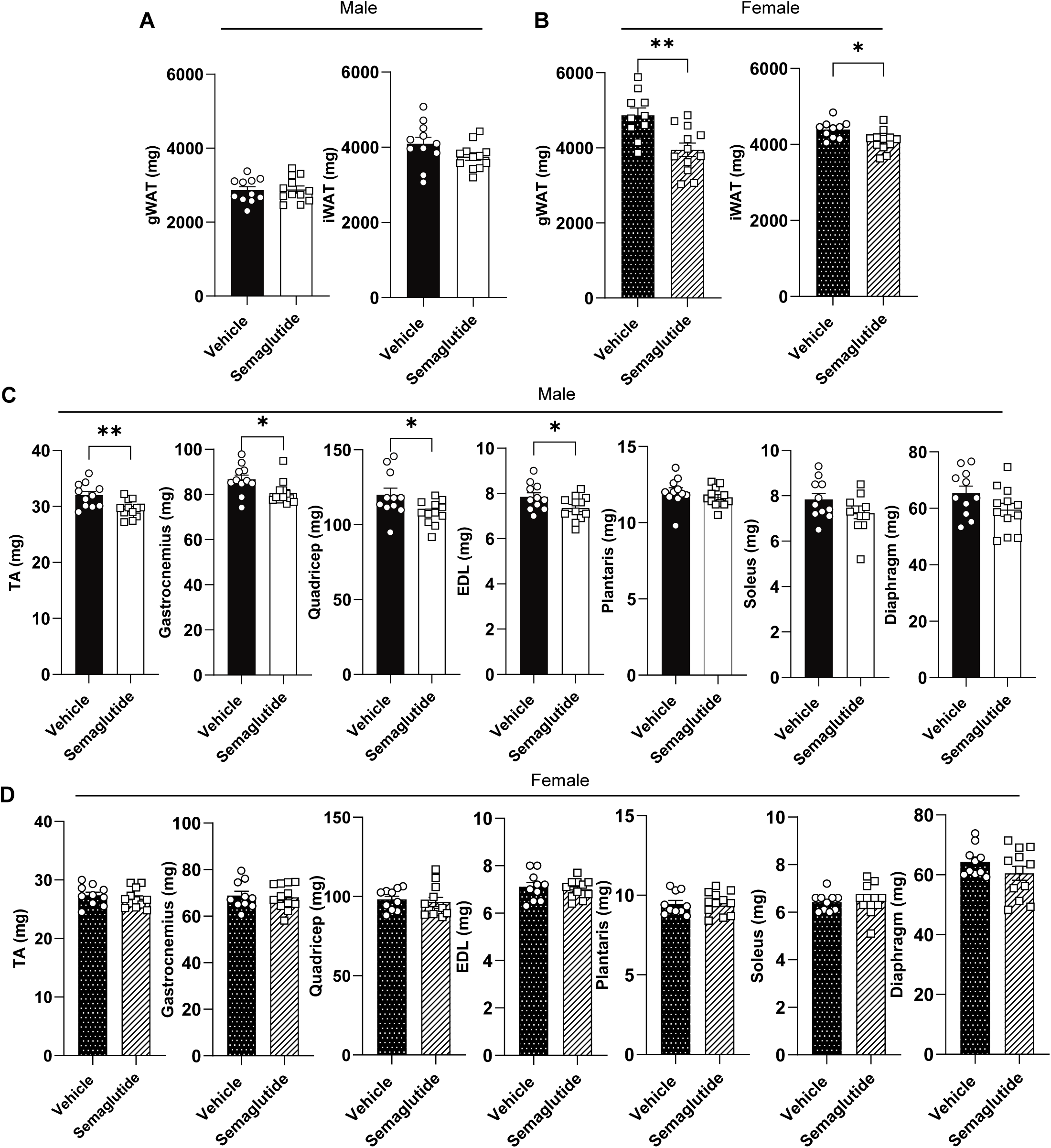
Semaglutide-induced sex-dependent remodeling of adipose and muscle mass. A: Showing gonadal white adipose tissue (gWAT) and inguinal white adipose tissue (iWAT) mass of vehicle or semaglutide treated male mice. B: gWAT and iWAT mass of vehicle or semaglutide treated female mice. C: Individual muscle mass of vehicle or semaglutide treated male ob/ob mice. D: Individual muscle mass of vehicle or semaglutide treated female *ob/ob* mice Data is represented as mean ± SEM. * p<0.05 and ** p<0.01 significant difference vs. corresponding controls. n=12/group for both male and female.

Sex-specific differences were also evident in skeletal muscle mass. Semaglutide-treated male *ob/ob* mice showed significant reductions in several muscles, including TA (∼7.6%) EDL (∼6.3%), gastrocnemius (∼6.8%), and quadriceps (∼9.8%) (Figure 2C, Table 1). In contrast, female mice did not exhibit significant reductions in any skeletal muscle compared with vehicle-treated controls (Figure 2D). Beyond skeletal muscle, semaglutide treatment reduced the mass of multiple lean tissues in both sexes. Apart from the lungs, the liver, spleen, heart, and kidneys were significantly reduced (Supplement figure 1A&B). Collectively, these findings reveal a sexually dimorphic response to semaglutide treatment in *ob/ob* mice, characterized by preferential adipose tissue mobilization in females and disproportionate reductions in skeletal muscle mass in males.

### Semaglutide treatment does not produce atrophy with no alteration in muscle force production in *ob/ob* mice

To determine whether the sex-specific difference in the lean mass alters skeletal muscle structure and fiber-type composition, we analyzed muscle fiber morphology and composition in the TA muscle. In males, semaglutide treatment did not alter mean fiber cross-sectional area (CSA) (Figure 3B) or the relative proportion of Type I, IIa, IIx, and IIb fibers compared with vehicle (Figure 3C). Females showed a modest but non-significant decrease in mean fiber CSA (Figure 3D) without changes in fiber type composition (Figure 3E).

**Figure 3.**
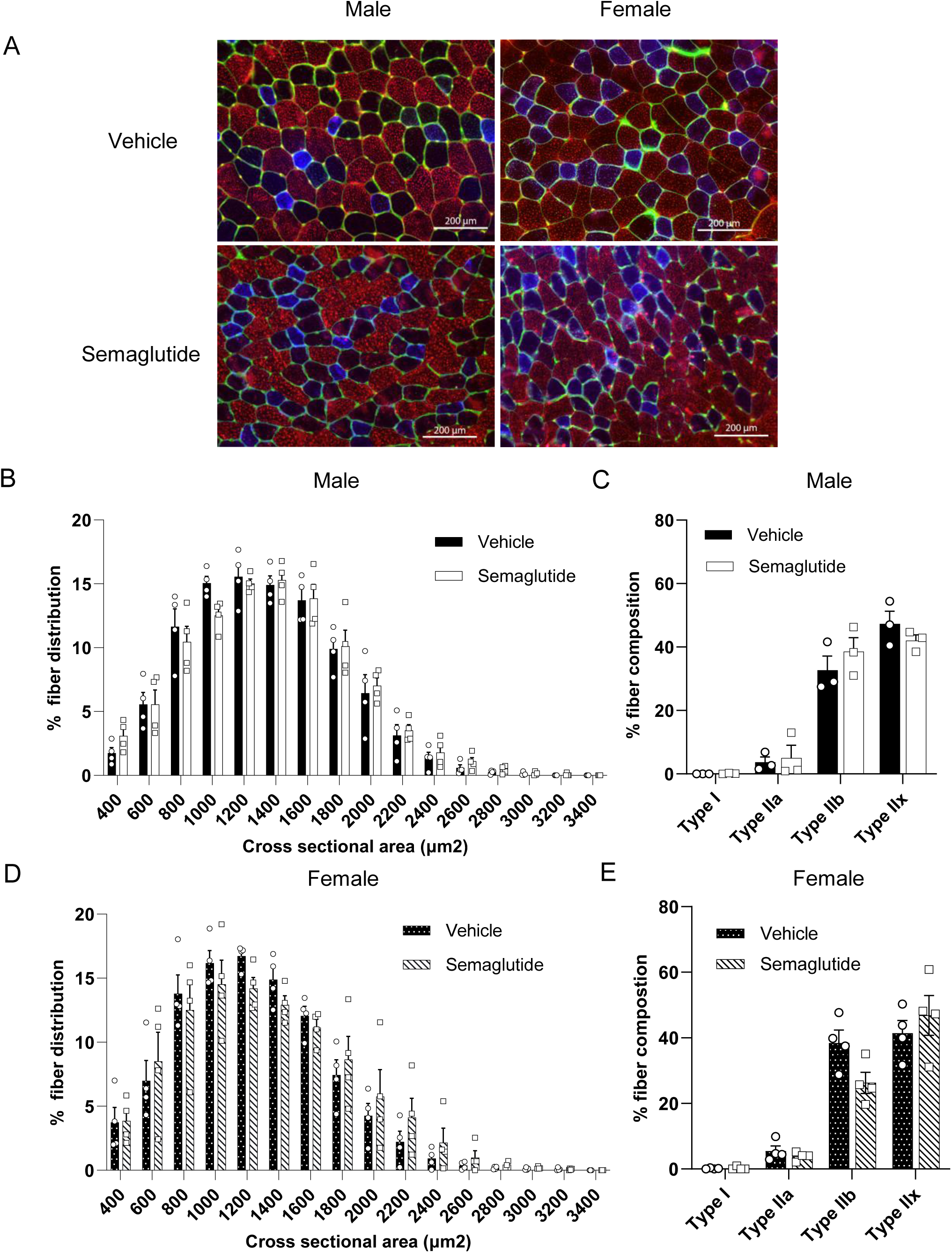
Semaglutide treatment did not alter muscle cross-sectional area or fiber types. A: A: Immunohistochemical characterization of tibialis anterior (TA) muscle fiber types. TA muscle cross-sections were stained to identify individual fiber types: Type IIb fibers appear red, Type IIa fibers appear blue, and unstained fibers (appearing black) correspond to Type IIx. Wheat germ agglutinin (WGA) labeling outlines muscle fiber borders in green, enabling clear visualization of fiber morphology and distribution. B & D: Showing muscle % fibers distribution per cross sectional area in vehicle or semaglutide treated male and female mice. C & E: Showing muscle % fiber composition in vehicle or semaglutide treated male and female mice. Data is represented as mean ± SEM. n=3/group for male and n=4/group for female.

We next examined whether semaglutide-induced muscle loss was associated with altered contractile function or muscle fiber morphology. The effect of three-week semaglutide treatment on muscle force production was assessed in both male and female *ob/ob* mice. Following treatment, EDL and soleus muscles were isolated to measure their force-producing capacity. Force-frequency analyses revealed no significant changes in the force production capacity of the soleus muscle in either males or females after semaglutide-induced weight loss. Consistent with these observations, the maximum tetanic force was also unaltered in the soleus of males or females (Figure 4A & B). Similarly, EDL muscles from both sexes showed no significant changes in force-frequency responses or maximal force generation following semaglutide treatment (Figure 4C & D).

**Figure 4.**
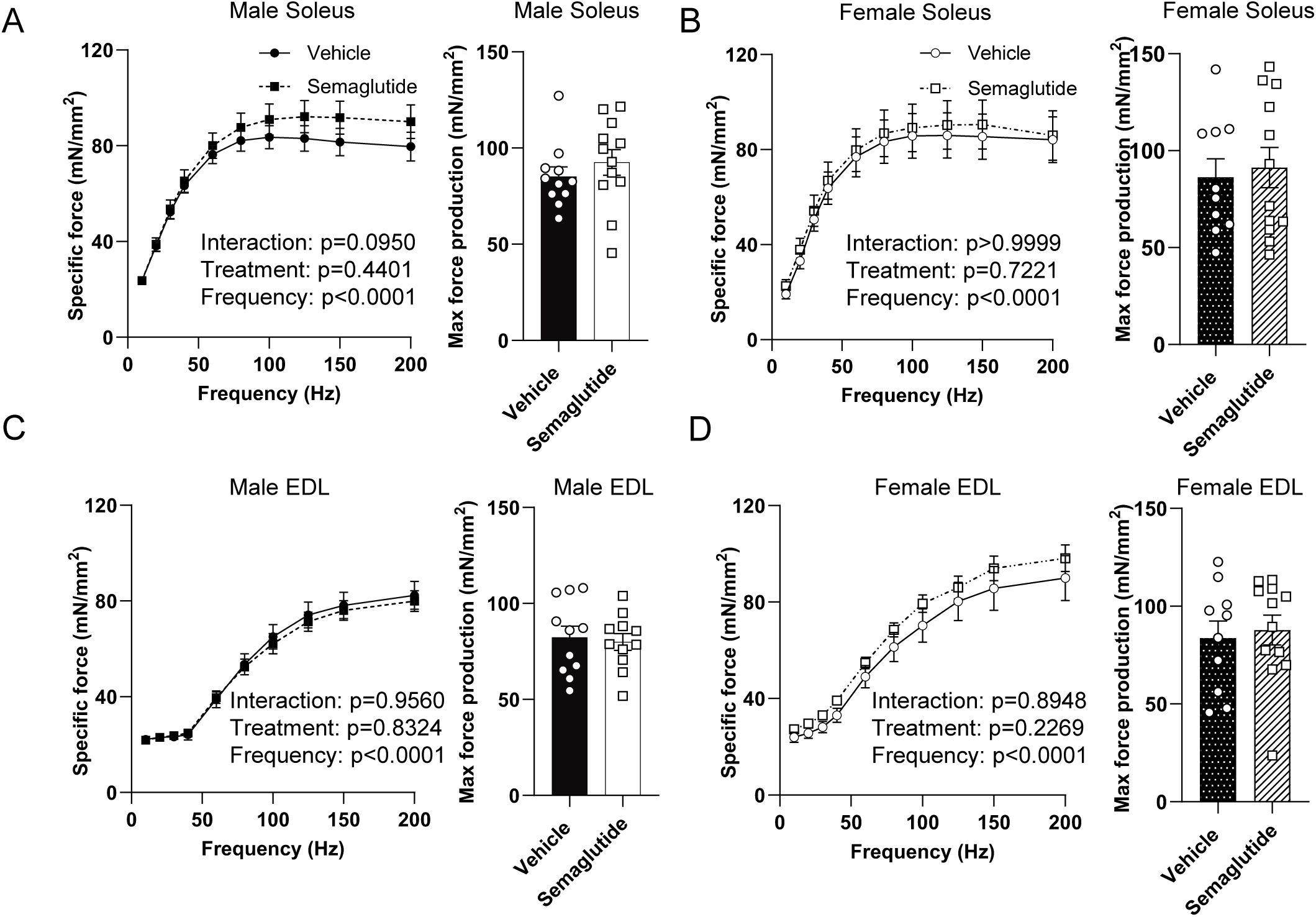
Semaglutide-induced weight loss did not affect the muscle force production capacity in both male and female *ob/ob* mice. A&B: The specific force and maximum force production capacity of soleus in both male and female mice in both the vehicle and semaglutide treated group. C&D: The specific force and maximum force production capacity of EDL between both the treatment groups in both the sexes. Data is represented as mean ± SEM. * p<0.05 significant difference vs. corresponding controls. n=12/group for both male and female.

### Semaglutide-induced weight loss does not improve mitochondrial OXPHOS efficiency in *ob/ob* mice

We previously demonstrated that weight loss from diet-induced obesity, whether achieved through semaglutide treatment or dietary intervention, improves skeletal muscle mitochondrial efficiency (15, 17) in C57BL6 mice. We therefore examined whether this adaptive response is preserved in leptin-deficient *ob/ob* mice. In contrast to our prior findings, semaglutide-induced weight loss did not significantly alter mitochondrial efficiency in either sex, as assessed by the P/O ratio (ATP produced per mole of O consumed) (Figure 5A). However, semaglutide treatment induced a transient increase in the rate of ATP production (*J*ATP) in male mice only, with no corresponding change observed in females (Supplement figure 3A). Importantly, semaglutide did not significantly affect mitochondrial oxygen consumption rates (*J*O) in either male or female *ob/ob* mice (Supplement figure 3B).

**Figure 5:**
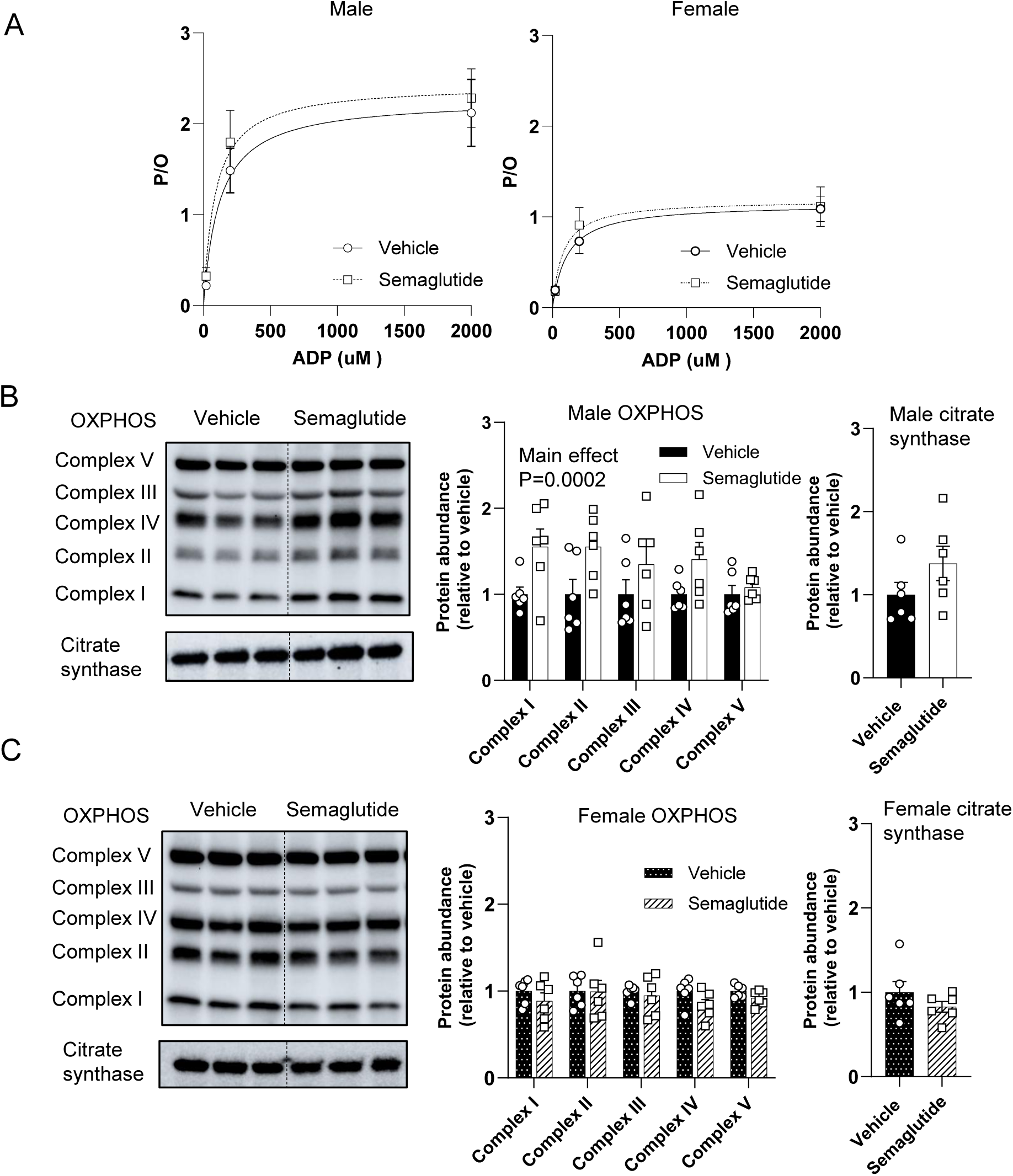
Semaglutide does improve skeletal muscle mitochondrial efficiency in *ob/ob* male mice. A: Mitochondrial efficiency (P/O ratio) of both vehicle and semaglutide treated male and female mice. B: Western blot analysis for mitochondrial OXPHOS subunits and citrate synthase for male mice. C: Western blot analysis for mitochondrial OXPHOS subunits and citrate synthase for female mice. Data is represented as mean ± SEM. * p<0.05 significant difference vs. corresponding controls. n=12/group in both male and female for high-respirometry analysis and n=6/group in both male and female for western blot.

To determine whether semaglutide-induced weight loss alters skeletal muscle mitochondrial content and oxidative capacity, we next assessed the abundance of mitochondrial OXPHOS complexes and citrate synthase activity. Male semaglutide-treated mice revealed significant increase in OXPHOS individual complex, while citrate synthase abundance remains unchanged (Figure 5B). However, female mice exhibited no detectable changes in both OXPHOS and citrate synthase abundance (Figure 5C). These findings indicate that semaglutide-induced weight loss does not alter skeletal muscle mitochondrial OXPHOS efficiency or content in leptin-deficient *ob/ob* mice.

## Discussion

Semaglutide reduces body weight primarily through suppression of appetite via POMC activation and AgRP inhibition (25–27). This neural loop normally operates in concert with leptin to regulate energy balance (28, 29). Given that leptin-deficient *ob/ob* mice develop severe obesity and systemic metabolic dysfunction across both sexes, they represent a powerful model for dissecting sex-specific adaptations to pharmacologically induced weight loss (30). In this study, we demonstrate that semaglutide-induced weight loss elicits pronounced sex-specific differences in body composition. Female *ob/ob* mice exhibited significant reductions in adipose tissue mass without concomitant loss of skeletal muscle, whereas male *ob/ob* mice showed a reduction in muscle mass with minimal changes in adipose tissue. These findings highlight a distinct sexual dimorphism in the tissue-specific effects of semaglutide.

A key finding of this study is the strong sexual dimorphism in tissue remodeling during semaglutide-induced weight loss. While both sexes achieved similar reductions in total body weight and body composition, individual tissues revealed distinct outcomes. Male *ob/ob* mice exhibited significant reductions in skeletal muscle mass without proportional decreases in visceral and subcutaneous adipose depots. In contrast, females preferentially decreased iWAT and gWAT while preserving skeletal muscle mass. This divergence indicates that while the gross efficacy of semaglutide is maintained across sexes, the underlying physiological processes differ. This sexual dimorphism may reflect intrinsic sex-specific protective mechanisms that preserve skeletal muscle during mild energy deficit. Although *ob/ob* mice are characterized by hypogonadism (31, 32), female-specific signaling pathways, such as residual estrogenic activity or XX-chromosomal factors, may still attenuate proteolysis. Estrogen is known to attenuate proteolysis through suppression of the FoxO3a/Atrogin-1/MuRF1 pathway and to enhance mitochondrial efficiency and oxidative metabolism (33). Therefore, under leptin deficiency, estrogen may provide a compensatory mechanism that counteracts semaglutide-induced muscle catabolism in females. Moreover, estrogen amplifies GLP-1R signaling in the hypothalamus, promoting stronger appetite suppression and greater caloric deficit in females (34, 35). Given that female mice typically possess a higher baseline fat mass due to estrogen-driven adiposity (36), the increased energy demand during semaglutide treatment is likely met through enhanced fat oxidation rather than muscle catabolism (37), thereby preserving lean tissue. Together, these mechanisms provide a plausible basis for the muscle-sparing phenotype observed in female *ob/ob* mice, although direct testing will be required to determine their specific contributions in the leptin-deficient state.

Given the fact that muscle contractile function is closely associated with fiber size and fiber-type composition (38–40), the preserved ex vivo force generation in semaglutide-treated mice is consistent with our histological findings. Despite modest reductions in hindlimb muscle mass in males, we observed no differences in force generation between vehicle- and semaglutide-treated groups of either sex. This aligns with the absence of changes in muscle fiber CSA or fiber-type distribution in the TA muscle. Fast-twitch fibers (Type IIa, IIx, and IIb) generally possess larger CSA and greater force-generating capacity compared with slow-twitch (Type I) fiber (38, 39, 41); thus, the preservation of this distribution likely contributed to the maintenance of muscle force. However, this mechanistic interpretation should be made cautiously, since force measurements were performed in the soleus and EDL muscles, whereas fiber typing was conducted in TA a predominantly fast-twitch muscle with minimal Type I fiber content. This pattern aligns with our previous work in DIO male mice using the same semaglutide dose, where we observed an approximately 20% reduction in EDL force after one week of the treatment that recovered by week three, despite no changes in fiber CSA or fiber-type composition (14). Thus, because the present study measured force-generating capacity only in week three, it is possible that early transient deficits in *ob/ob* mice had already resolved before measurement.

Weight loss induced by dietary intervention or semaglutide treatment in DIO mice has been shown to improve OXPHOS efficiency in skeletal muscle, reflected by an increased P/O ratio which represents the amount of ATP produced per unit of O_2_ consumed (17). To determine whether this adaptive response occurs in leptin-deficient *ob/ob* mice, we assessed mitochondrial respiration and ATP production after three weeks of semaglutide treatment. In contrast to findings in DIO mice, semaglutide did not significantly alter *J*O_2_, *J*ATP, or the resulting P/O ratio in either sex, indicating no improvement in mitochondrial efficiency. However, mitochondrial complex abundance was improved in male mice while citrate synthases activity did not differ between semaglutide- and vehicle-treated groups. These results suggest that the improved mitochondrial efficiency typically observed with weight loss in leptin-intact models does not occur in the absence of leptin signaling. Leptin deficiency has been shown to impair AMPK-PGC1α signaling in skeletal muscle (42) and promote lipid overload, incomplete β-oxidation, and elevated mitochondrial reducing pressure conditions that may blunt the improvements in OXPHOS coupling efficiency typically seen with weight loss (43, 44). Together, these findings indicate that leptin signaling is a critical permissive factor for mitochondrial efficiency improvements during semaglutide-induced weight loss.

In summary, semaglutide induced modest weight loss in leptin-deficient *ob/ob* mice but produced marked sexual dimorphism in tissue remodeling. Female mice exhibited preferential adipose tissue loss with preservation of skeletal muscle mass, accompanied by a rightward shift in fiber CSA distribution and a transition from Type IIb to Type IIx fibers, consistent with fast-twitch remodeling without functional compromise. In contrast, male mice demonstrated a reduction in skeletal muscle mass with a selective decrease in Type IIx fiber proportion, despite preservation of overall fiber CSA distribution. Importantly, muscle force-generating capacity remained intact in both sexes, indicating that the observed sex-specific alterations in fiber composition did not translate into impaired contractile function at the three-week time point. Although leptin deficiency does not fully recapitulate common human obesity, these findings highlight pronounced sex-specific differences in skeletal muscle and adipose adaptations to semaglutide-induced weight loss and underscore the importance of considering biological sex when evaluating the metabolic and body composition effects of GLP-1 receptor agonist therapies

## Acknowledgment

This research is supported by NIH grants (DK107397, DK127979, GM144613, and AG074535 to K.F., CA278826 to K.H.F, CA286584 and AG065993 to A.H.C), and the Grant-in-aid for Japan, Society for Promotion of Science (JSPS) Fellows (24KJ2039 to T.K.). University of Utah, Metabolomics Core Facility is supported by DK110858.

The authors declared no conflict of interest.

S.R, K.F, R.H.C, A.C and M.J.D designed the study. S.R, T.K, and R.H.C performed the mouse, muscle force and mitochondrial experiment and analyzed data. S.W analyzed muscle fibers from histology. S.R, K.F and R.H.C wrote the manuscript with edits from all the authors.

The authors received no external writing assistance in the preparation of this manuscript. Editorial and scientific input were provided solely by the listed authors.

Dr. Ran Hee Choi and Dr. Katsuhiko Funai are the guarantors of this work and, as such, had full access to all the data in the study and took responsibility for the integrity of the data and the accuracy of the data analysis.

Prior presentation in abstract form in the conference Obesity week, November 4, 2025.

## Supplement Figures

**Supplement Figure 1.**
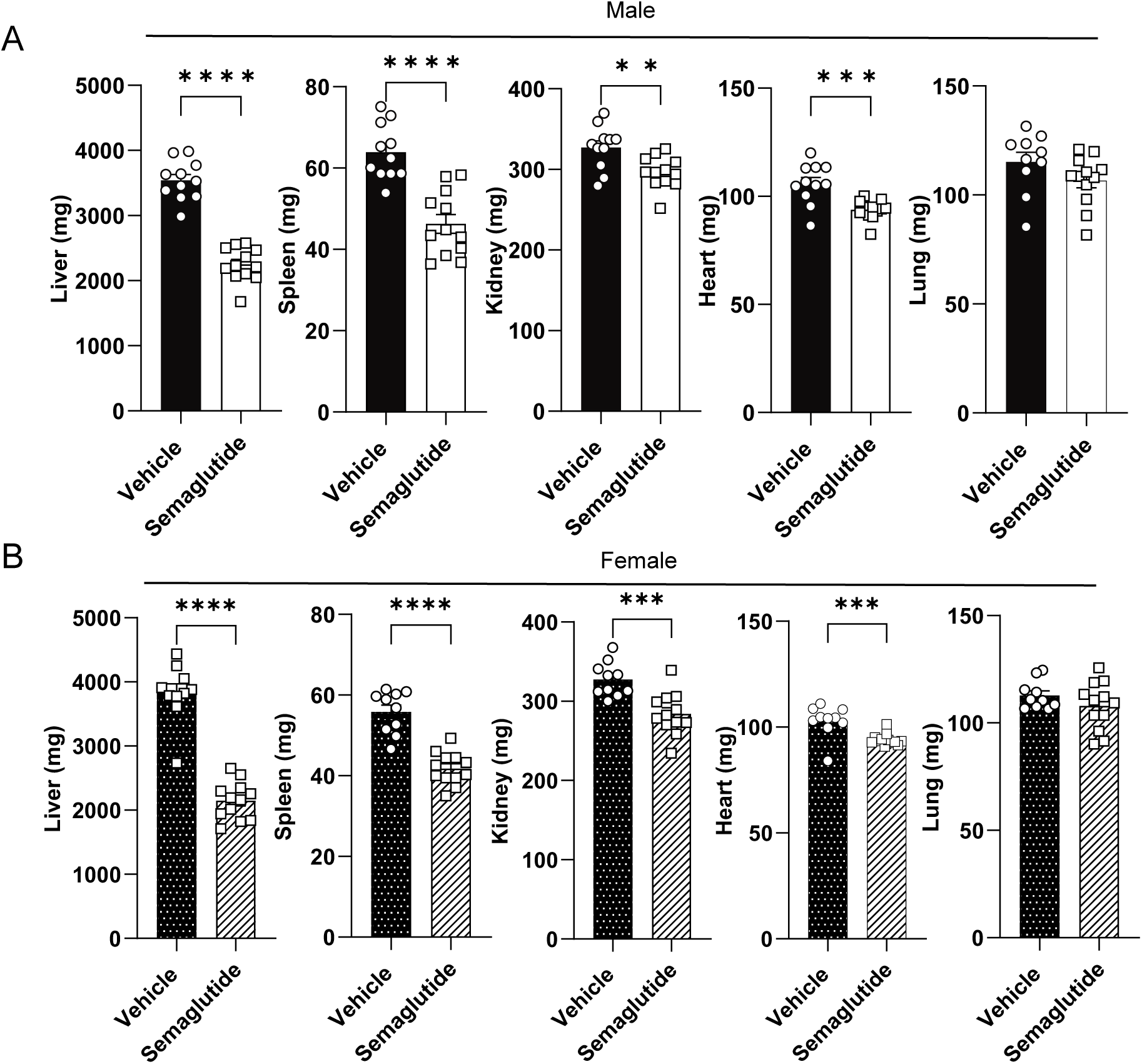
Semaglutide induced weight loss promotes lean tissue mass reduction. A: Individual mass of liver, spleen, kidney, heart masses in vehicle or semaglutide treated male mice. B: Individual lean tissue mass of vehicle and semaglutide treated female mice. Data is represented as mean ± SEM. ** p<0.01, *** p<0.001 and **** p<0.0001 significant difference vs. corresponding controls. n=12/group for both male and female.

**Supplement Figure 2.**
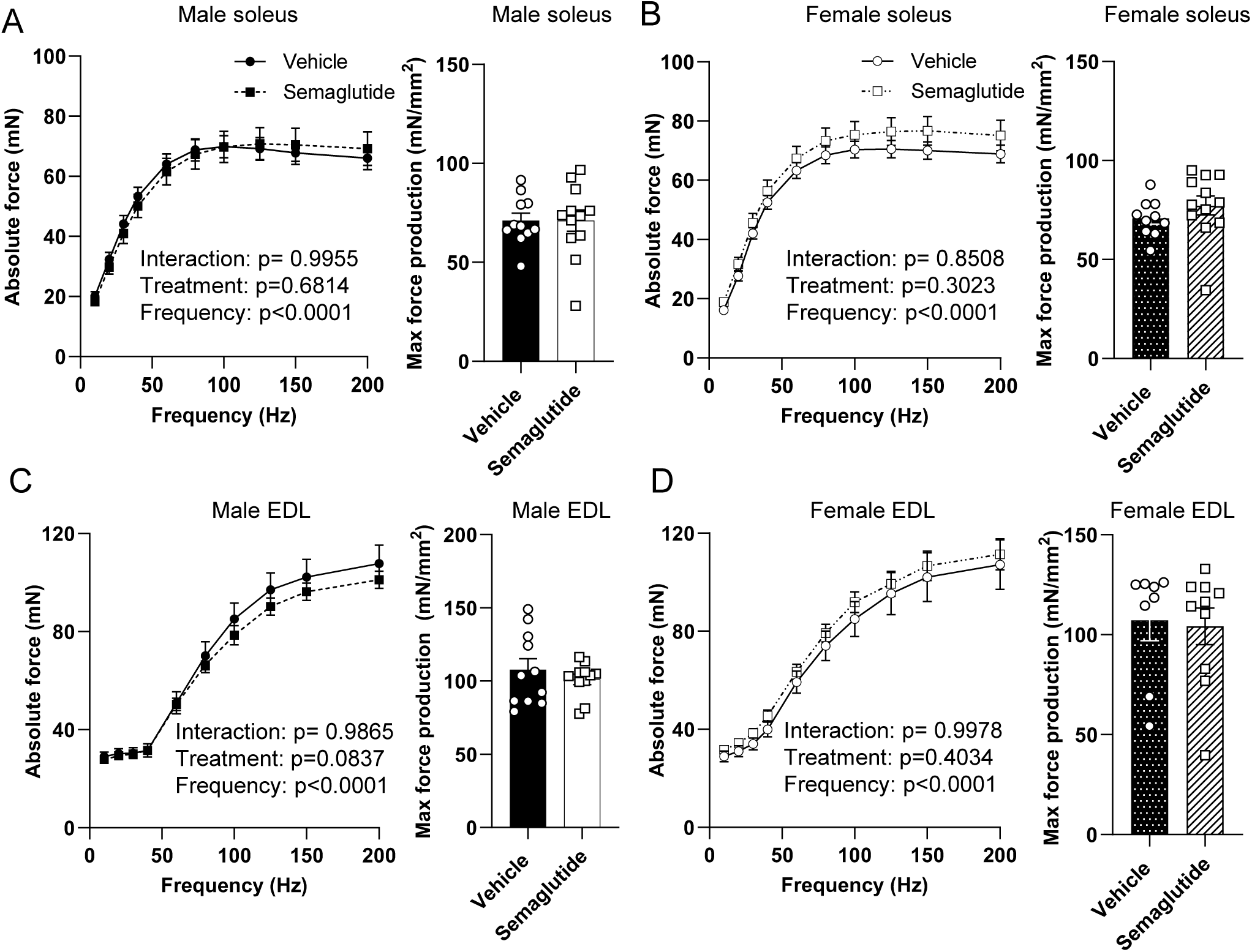
No alteration in the absolute force production capacity with semaglutide treatment. A&B: The absolute force production capacity of soleus in both male and female semaglutide treated mice compared to vehicle treated group. C&D: The absolute force production capacity of EDL in vehicle and semaglutide treated in both the sexes. Data is represented as mean ± SEM. * p<0.05 significant difference vs. corresponding controls. n=12/group for both male and female.

**Supplement Figure 3.**
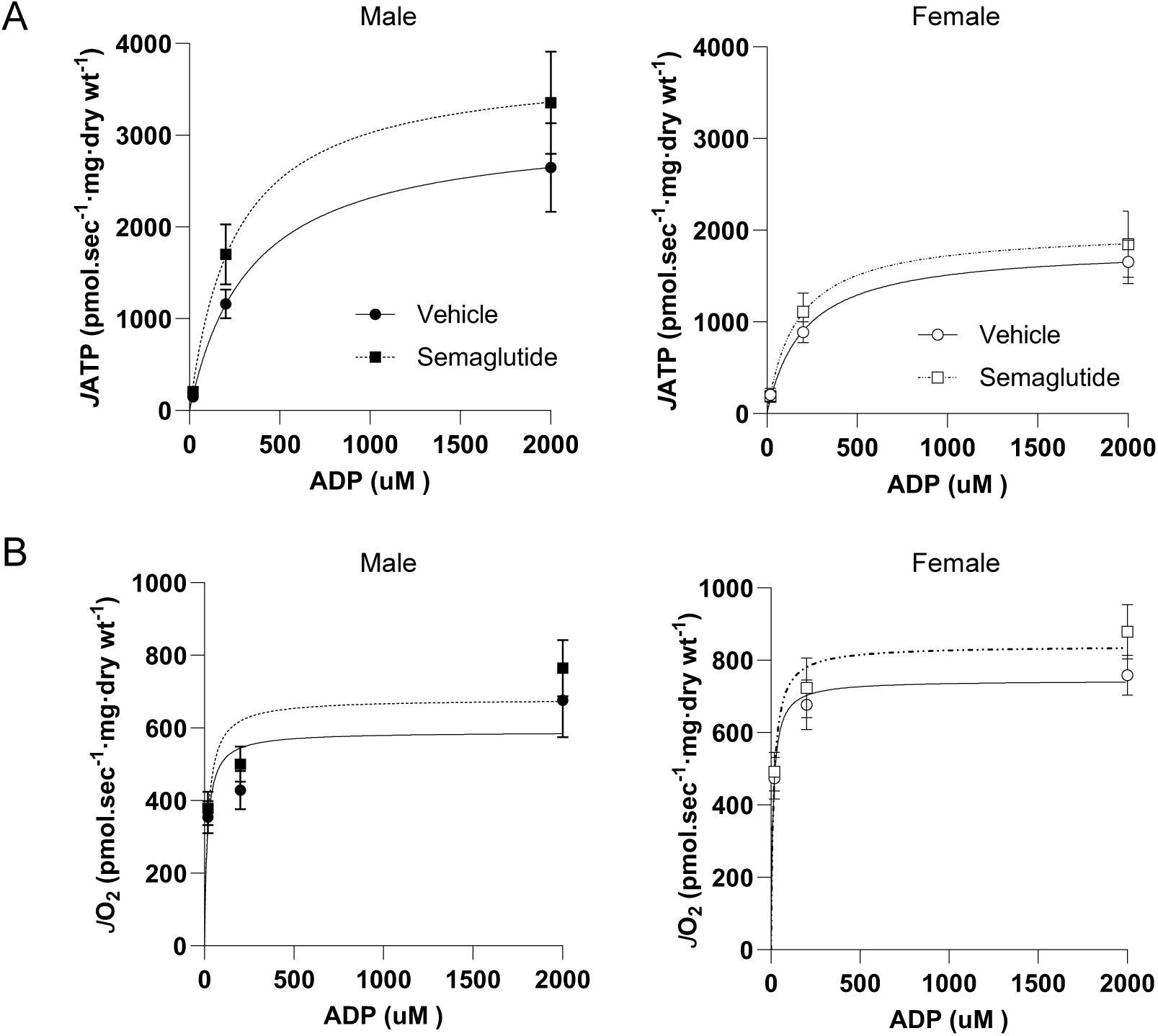
Effects of semaglutide on skeletal muscle mitochondrial bioenergetics in *ob/ob* mice. A: The rate of ATP production (*J*ATP) in semaglutide or vehicle treated male and female *ob/ob* mice. B: The rate of oxygen consumption (*J*O_2_) in semaglutide or vehicle treated male and female *ob/ob* mice. Data is represented as mean ± SEM. * p<0.05 significant difference vs. corresponding controls. n=12/group in both male and female for high-respirometry analysis.

